# Predictive visual motion extrapolation emerges spontaneously and without supervision at each layer of a hierarchical neural network with spike-timing-dependent plasticity

**DOI:** 10.1101/2020.08.01.232595

**Authors:** Anthony N. Burkitt, Hinze Hogendoorn

## Abstract

The fact that the transmission and processing of visual information in the brain takes time presents a problem for the accurate real-time localisation of a moving object. One way this problem might be solved is extrapolation: using an object’s past trajectory to predict its location in the present moment. Here, we investigate how a simulated *in silico* layered neural network might implement such extrapolation mechanisms, and how the necessary neural circuits might develop. We allowed an unsupervised hierarchical network of velocity-tuned neurons to learn its connectivity through spike-timing dependent plasticity. We show that the temporal contingencies between the different neural populations that are activated by an object as it moves causes the receptive fields of higher-level neurons to shift in the direction opposite to their preferred direction of motion. The result is that neural populations spontaneously start to represent moving objects as being further along their trajectory than where they were physically detected. Due to the inherent delays of neural transmission, this effectively compensates for (part of) those delays by bringing the represented position of a moving object closer to its instantaneous position in the world. Finally, we show that this model accurately predicts the pattern of perceptual mislocalisation that arises when human observers are required to localise a moving object relative to a flashed static object (the flash-lag effect).

**Significance Statement:** Our ability to track and respond to rapidly changing visual stimuli, such as a fast moving tennis ball, indicates that the brain is capable of extrapolating the trajectory of a moving object in order to predict its current position, despite the delays that result from neural transmission. Here we show how the neural circuits underlying this ability can be learned through spike-timing dependent synaptic plasticity, and that these circuits emerge spontaneously and without supervision. This demonstrates how the neural transmission delays can, in part, be compensated to implement the extrapolation mechanisms required to predict where a moving object is at the present moment.

## 1. Introduction

The transmission and processing of information in the nervous system takes time. In the case of visual input to the eyes, for example, it takes up to ~50-70 milliseconds for information from the retina to reach the primary visual cortex (Maunsell & Gibson, 1992; Lamme & Roelfsema, 2000), and up to ~120 milliseconds before human observers are able to initiate the first actions based on that information (Thorpe et al., 1996; Kirchner & Thorpe, 2006). Because events in the world continue to unfold during this time, visual information becomes progressively outdated as it travels up the visual hierarchy.

Although this is inconsequential when visual stimuli are unchanging on this time scale, these delays pose a problem when input is time-varying, for instance in the case of visual motion. If neural delays were not somehow compensated, we would consistently mislocalise moving objects behind their true positions. However, humans and many other visual animals are strikingly accurate at interacting with even fast moving objects (Smeets et al., 1998), suggesting that the brain implements some kind of mechanism to compensate for neural delays.

One candidate mechanism by which the brain might compensate for delays is prediction (Nijhawan, 2008). In the case of motion, the brain might use an object’s previous trajectory to extrapolate its current position, even though actual sensory input about the object’s current position will not become available for some time. Consistent with this interpretation, motion extrapolation mechanisms have been demonstrated in multiple levels of the visual hierarchy, including the retina of salamanders, mice, and rabbits (Berry et al., 1999; Hosoya et al., 2005; Schwartz et al., 2007), cat lateral geniculate nucleus (Sillito et al., 1994), and both cat and macaque primary visual cortex (Jancke et al., 2004; Subramaniyan et al., 2018; Benvenuti et al., 2019). In humans, recent EEG and MEG studies using apparent motion similarly revealed predictive activation along motion trajectories (Hogendoorn & Burkitt, 2018; Aitken et al., 2020; Blom et al., 2020; Robinson et al., 2020), and motion extrapolation mechanisms have been argued to be the cause of the so-called flash-lag effect (Hogendoorn, 2020; Nijhawan, 1994; Khoei et al., 2017).

The existence of predictive mechanisms at multiple stages of the visual hierarchy is reminiscent of hierarchical predictive coding, a highly influential model of cortical organisation (Rao & Ballard, 1999). In this model, multiple layers of a sensory hierarchy send predictions down to lower levels, which in turn send prediction errors up to higher levels. In this way, the hierarchy essentially infers the underlying causes of incoming sensory input, using prediction errors to correct and update that inference. It is important to note, however, that the “predictions” in predictive coding are hierarchical, rather than temporal: predictive coding networks ‘predict’ (or reconstruct) activity patterns in other layers, rather than predicting the future. Consequently, the conventional formulation of predictive coding cannot compensate for neural delays. In fact, we previously argued that neural delays pose a specific problem for hierarchical predictive coding, because descending hierarchical predictions will be misaligned in time with ascending sensory input (Hogendoorn and Burkitt, 2019). For any time-varying stimulus (such as a moving object) this would lead to significant (and undesirable) prediction errors.

To address this, we previously proposed a Real-Time Temporal Alignment hypothesis, which extends the predictive coding framework to account for neural transmission delays (Hogendoorn and Burkitt, 2019). In this hypothesis, both forward and backward connections between hierarchical layers implement extrapolation mechanisms to compensate for the incremental delay incurred at that particular step. Without these extrapolation mechanisms, delays progressively accumulate as information flows through the visual hierarchy, such that information at higher hierarchical layers is outdated relative to information at lower hierarchical layers. Conversely, the striking consequence of the Real-Time Temporal Alignment hypothesis is that for a predictable stimulus trajectory, different layers of the visual hierarchy become aligned in time. The hypothesis posits that extrapolation mechanisms are implemented at multiple stages of the visual system, which is consistent with the neurophysiological findings outlined above, as well as with human behavioural experiments (van Heusden et al., 2019). However, a key question that remains is how such extrapolation mechanisms are implemented at the circuit level, and how those neural circuits arise during development.

Here, we address those two questions by simulating *in silico* the first two layers of a feedforward hierarchical neural network sensitive to visual motion. We present the network with simulated moving objects, and allow neurons to learn their connections through spike-time dependent plasticity (STDP; Markram et al., 1997; Bi & Poo, 1998), a synaptic learning rule that strengthens and weakens synapses contingent on the relative timing of input and output action potentials. We focus on the first two layers of the hierarchical network to explore the key mechanisms, which would be expected to occur at each higher level of the hierarchy.

We show that when a motion-sensitive hierarchical network is allowed to learn its connectivity through STDP (without supervision), the temporal contingencies between the different neural populations that are activated by the object as it moves cause the receptive fields of higher-level neurons to spontaneously shift in the direction opposite to their preferred direction of motion. As a result, they start to encode the extrapolated position of a moving object along its trajectory, rather than its physical position. However, due to the delays inherent in neural transmission, this mechanism actually brings the represented position of the object closer to its instantaneous position in the world, effectively compensating for (part of) those delays. Finally, we show that the behaviour of the resulting network predicts the pattern of velocity-dependence in the perceptual localisation of moving objects.

## 2. Methods

### 2.1. Network Architecture

The network architecture considered here is one in which a moving object generates a visual input stimulus that is encoded at each layer of the network by a population code that represents both the position and velocity of the stimulus. This population code includes sub-populations of neurons tuned to both position and velocity, as has previously been proposed by Khoei et al. (2017) and consistent with the known velocity-tuning of a large proportion of visual neurons in the early visual system (Orban et al., 1986). This sub-population coding of the velocity of the stimulus at each layer is inherited from the lower layers, beginning in the retina (Berry et al., 1999; Ravello et al, 2019) and LGN (Sillito et al., 1994), and then passed on to the primary visual cortex (Janke et al., 2004; Benvenuti et al., 2020), and it is further enhanced in the motion-processing centers of areas MT and MST (Maunsell & Van Essen, 1983; Koch et al., 1989; Perrone & Thiele, 2001; Inaba et al., 2011).

Three layers of the network are shown schematically in Figure 1A, in which there are *N*_n_ position sub-populations at each layer. Although both classical predictive coding and the Real-Time Temporal Alignment hypothesis posit both feed-forward and feedback connections, here we consider only feed-forward connections as proof-of-principle. In addition, in a more general scheme lateral weights at each layer could also be included, as previously proposed by Jancke and Erlhagen (Jancke & Erlhagen, 2010), but these are neglected here.

**Figure 1:**
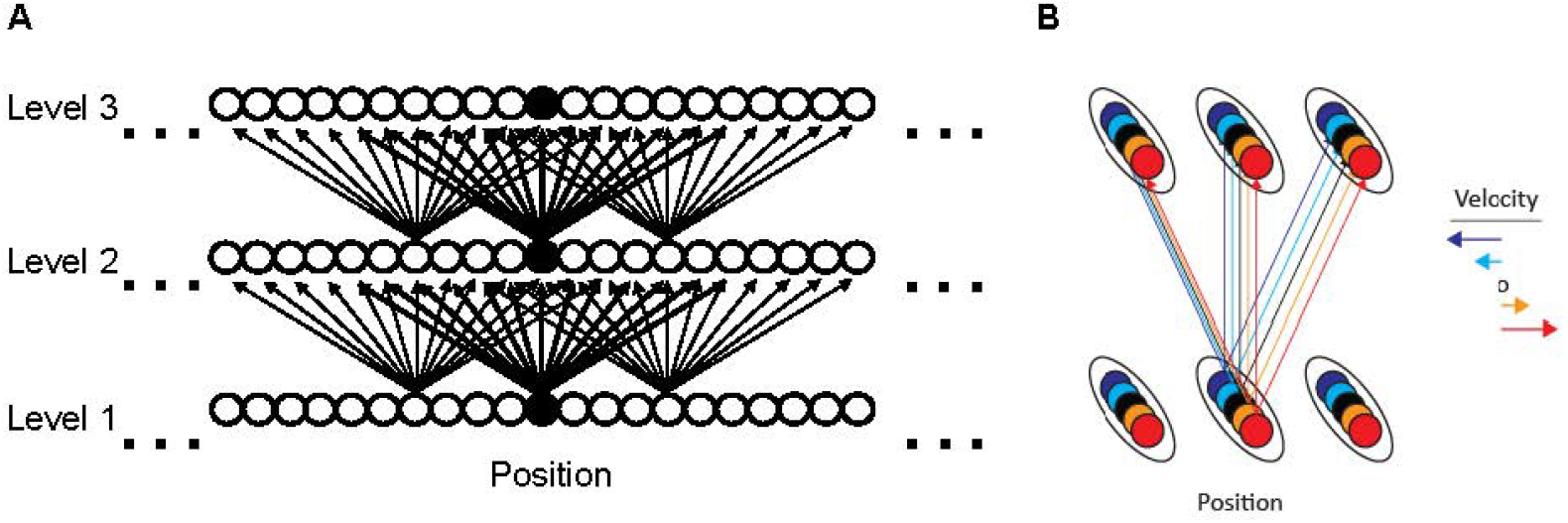
A. Schematic illustration of a portion of the first three layers of the hierarchical network architecture. The circles denote the neural population at each of the N_n_ positions (fifteen shown, with the central population in black). The straight lines indicate the possible non-zero weights connecting the neural populations between layers, which have a limited spatial spread. The arrows indicate the direction of the neural connectivity (pre-synaptic to post-synaptic) and only the weights for a subset of neurons is illustrated. B. Each position population is further divided into M velocity-tuned sub-populations. Each velocity-tuned sub-population projects to sub-populations in the subsequent layer with the same velocity-tuning. Note that the analysis in this study focusses upon two adjacent layers.

The neural activation of each stage of processing feeds forward to the following stage. An important aspect of the network architecture is that each neural population receives input from a limited receptive field of neural populations at the preceding stage. In this way, the receptive field size of neural populations increases as the activity propagates to higher stages of processing.

### 2.2 Neural Model

The Poisson neuron model is used, in which the spikes of each neuron *i* (*i* = *1*,…,*N*) are generated stochastically with a spiking-rate function, *r_i_*(*t*), that is time-dependent and described by a Poisson point process. The instantaneous probability of a spike is given by this Poisson spiking-rate function, *r_i_*(*t*), so that in numerical simulations with discrete time-steps of *Δt* the probability of neuron *i* firing a spike at time *t* is *r_i_*(*t*)*Δt*. This is a stochastic neuron model that is widely used for both analytical and computational studies (Gerstner et al., 2014 [Ch.7]). A more complete mathematical description and analysis of this model is given in Appendix A of (Kempter et al., 1998).

The population place code of the stimulus at every stage of processing is described by a set of *N*_n_ units representing overlapping place fields, equally distributed over the interval [0, 1] and each with an identical Gaussian distribution width *σ*_p_. This width represents the number of *independent* place fields, *N*_p_, in the input layer, given by *N*_p_ = 1/ *σ*_p_. In this network, the Gaussian distribution represents the activity evoked by a stimulus, as illustrated in Figure 2, in which the neural activity of each population of active input neurons corresponding to a particular place field (i.e., an object at a particular location) is represented by a different colour, and the Gaussian curve represents the amplitude of the firing-rate of each neuron in that population. The firing rate, which represents the rate of action potentials, is described by a Poisson process, with a base firing rate of 5Hz in the absence of stimulation. Note that periodic boundary conditions are used, so that the position code can equivalently be represented by place on a circle. For simplicity, we consider the situation where only one object activates the input at any time, so that the relative activations of the input neurons give a neural representation of the position of the object.

**Figure 2:**
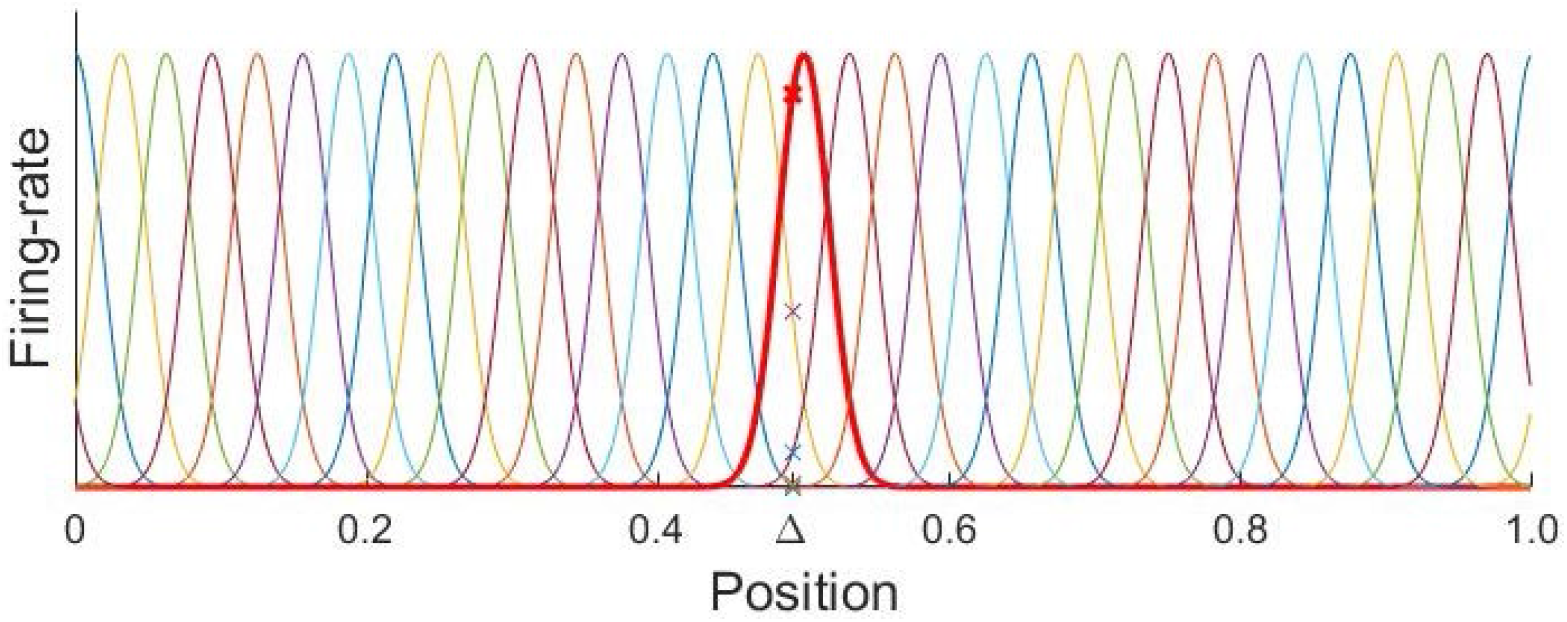
Place code in first (input) layer: Each colour represents the neural activity of the population of neurons, illustrated here for N_p_ = 32 positions corresponding to the location of the peak of each Gaussian curve, with width σ_p_ = 1/N_p_. A stimulus at the position indicated on the plot by Δ generates firing rates in the neural population, whose amplitude is indicated by the crosses on the corresponding place curves. The place distribution of population activity for a stimulus centred at the position x = 0.5 is shown in bold-red.

A layered network structure is considered, in which the units at the input layer feed their activity forward to the following layer, which has the same number of units, *N*_n_. For simplicity, the layers are taken to have an equal spatial separation and the propagation delay time for activity between layers has a constant value *t*_delay_. This neural transmission delay is of the order of several milliseconds between layers of the hierarchy (Maunsell & Gibson, 1992).

To incorporate velocity, each place field is further subdivided into *M* distinct sub-populations, corresponding to *M* different velocities (or velocity intervals) for the input stimulus, as illustrated in Figure 1B. The velocity of the object is encoded by the activity in the corresponding sub-population of the place fields. Consequently, an object moving at a constant velocity will primarily activate one velocity sub-population within each place field, i.e., for simplicity a discrete (quantised) representation of velocity is considered rather than a continuous representation. The velocity is assumed here to have been encoded by an earlier stage of neural processing, such as in the retina, so that the details of how this encoding occurs are not incorporated into the model. It suffices that velocity is encoded at each stage of the processing, which is a reasonable assumption as it is known that velocity is present in higher stages of the visual pathway such as area MT (Movshon & Newsome, 1996). In addition, the velocity sub-populations are assumed for simplicity to be precise, with no diffusion in velocity space. A more complete description of this process would include lateral connections between the neurons in each layer (both across positions and velocities, as implemented in Khoei et al. (2017) and Khoei et al. (2013)). However, the dynamics and interactions of these lateral connections are not the focus of this paper, since we are concerned here with the first wave of feed-forward activity through the network. Note that the place field at time *t* is defined as the average marginalized over the velocity sub-populations: *P*(*x*(*t*)) = ∑_*v*_ *P*(*x*(*t*)|*v*).

### 2.3 Neural Learning

We require that the computations that underlie learning in the network must be based upon known principles of synaptic plasticity, namely that the change in a synaptic strength is activity dependent and local. The locality constraint of synaptic plasticity requires that changes in the synaptic strength (i.e., the weight of a connection) can only depend on the activity of the presynaptic neuron and the postsynaptic neuron. Consequently, the spatial distribution of the synaptic changes in response to a stimulus are confined to the spatial extent of the position representation of the stimulus, which has important consequences for the structure of the network that emerges as a result of learning.

In the full network the weights are described by a matrix, ***W*** between every pair of successive layers and these are taken to be excitatory, in keeping with the excitatory nature of the long-range pyramidal neurons in cortex. Since our focus here will be upon the first two layers, ***W*** is taken to be a *N_n_*x*N_n_* matrix in which the elements *w*_ji_ are the weights between the first two layers. The locality constraint is implemented in the network by requiring that the weights from a neuron at location *i* in the first layer has a probability of being connected to a neuron at location *j* in the second layer that is Gaussian, namely that

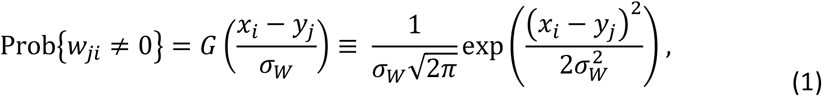

where *σ*_W_ is the width of the Gaussian distribution, *G*(·), and *x_i_* and *y_j_* are the locations of the neurons *i* and *j*, in the first- and second-layers resp. The width *σ*_W_ is chosen throughout to be sufficiently small relative to the spatial extent of the network (i.e., *σ*_W_ ≪ 1) that it is effectively equivalent to the van Mises circular distribution. In simulations, the amplitudes of the non-zero weights are initialized randomly and given small positive initial values, *w*_init_ = 0.01, while the zero-valued weights are fixed throughout (i.e., corresponding to the absence of any synaptic connection between the two neurons). In simulations this Gaussian distribution was truncated to zero at 5*σ*_W_. The non-zero weights then evolve according to Spike-Timing Dependent Plasticity (STDP), namely a change of the weight, Δ*w_ji_*, will occur when an input spike generated by neuron *i* at time *t_i_* arrives after a time-delay *t*_delay_ at a synapse on neuron *j*, and an output spike occurs at this neuron at time *t_j_*, so that the time difference Δ*t* lies within the STDP time-window *F_STDP_*(Δ*t*)

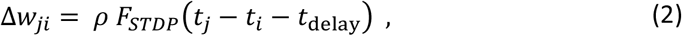

where *ρ* is the learning rate and

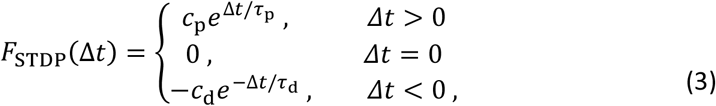

with coefficients of potentiation and depression given by *c*_p_ and *c*_d_, and time constants of potentiation and depression given by *τ*_p_ and *τ*_d_ resp., and Δ*t* = (*t_j_* − *t_i_* − *t*_delay_) is the difference between the spike output of the post-synaptic neuron at time *t_j_* and the incoming spike (generated by neuron *i* at time *t_i_*) that arrives at the synapse at time *t_i_* + *t*_delay_. Throughout the simulations balanced STDP was used, i.e., the STDP time-window has an equal amount of potentiation and depression, with *c*_p_ = *c*_d_ = 1 and *τ*_p_ = *τ*_d_ = 20ms, and a value of *t*_delay_ = 20 ms was chosen. An upper bound on the individual weights, *w*_max_, ensures that the weights do not grow unbounded. Note that the STDP learning described here is translationally invariant, since each neuron in a layer of the hierarchy receives a time- and space-shifted version of the input received by other neurons in the layer. Consequently, the learning is convolutional, because the weights connecting a neuron to neurons in the preceding layer of the hierarchy evolve to have a translational invariance across the layer.

Consider how an arbitrary weight *w_ij_* between unit *i* in Layer 1 at position *x_i_* and unit *j* in Layer 2 at position *y_j_* (chosen here for simplicity to be *y_j_* = 0) evolves according to STDP due the motion of an input stimulus moving from left to right with velocity *v*. Initially each weight has a small value *w_0_* and the distribution of possible weights is given by Eq.(1). Suppose that the stimulus of unit *i* is given by *r_i_*(*t*), then the change in weight *Δw_ji_* will depend upon both the timing of the pre-synaptic (input) spikes, with probability distribution *P*(*t_i_*) ∝ *r_i_*(*t*), and the timing of the post-synaptic (output) spikes, with probability distribution *P*(*t_j_*):

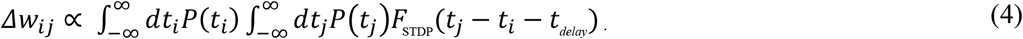

Consider now a stimulus that is a very localized point-like object, such as an insect or the spot or light from a laser pointer or a dot in a circular motion on a screen, moving at velocity *v*. This input can be approximated as a Dirac delta function, 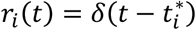, where 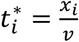. Then the probability distribution *P*(*t_j_*) becomes

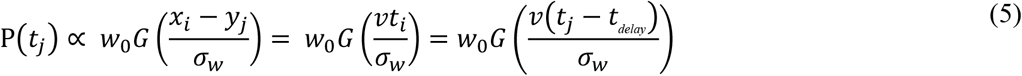

where *G*(·) is the Gaussian distribution given in Eq.(1). Consequently

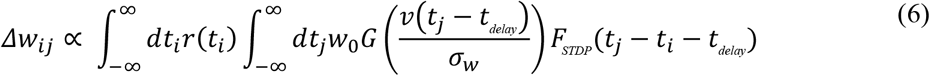

The time integral can be separated into the potentiation and depression contributions as

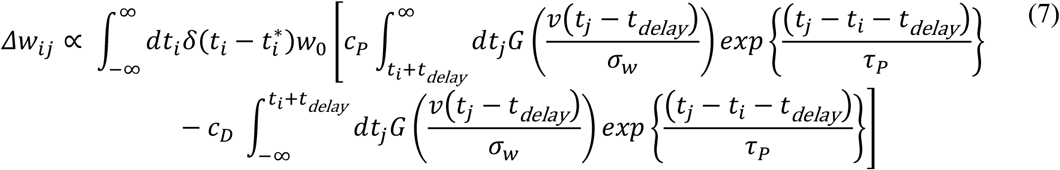

Integrating over *t_i_* using the properties of the Dirac delta function gives

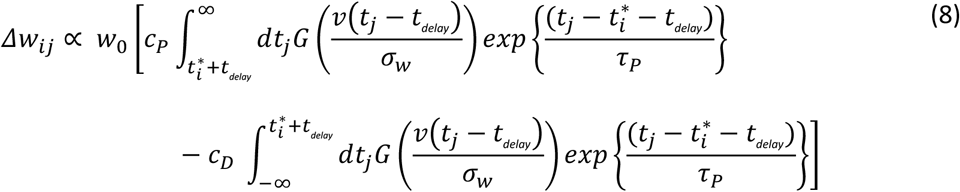

Assuming STDP with equal potentiation and depression *(τ* = *τ_P_* = *τ_D_* and *c* = *c_P_* = *c_D_*) gives

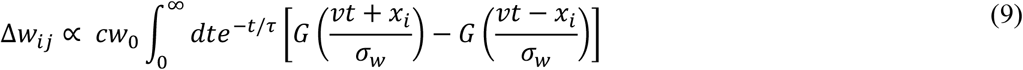

This change in the weight *w_ij_* as a function of the position *x_i_* of the input neuron is plotted in Figure 3 for velocities *v* = 0.05, 0.1, 0.15, 0.2 cycles/sec. The weights on the leading (incoming) side of the weight distribution experience more potentiation than depression due to STDP, whereas the weights on the tailing (outgoing) side of the weight distribution experience more depression than potentiation. This effect becomes larger as the velocity increases and the relative change in weights becomes larger.

**Figure 3:**
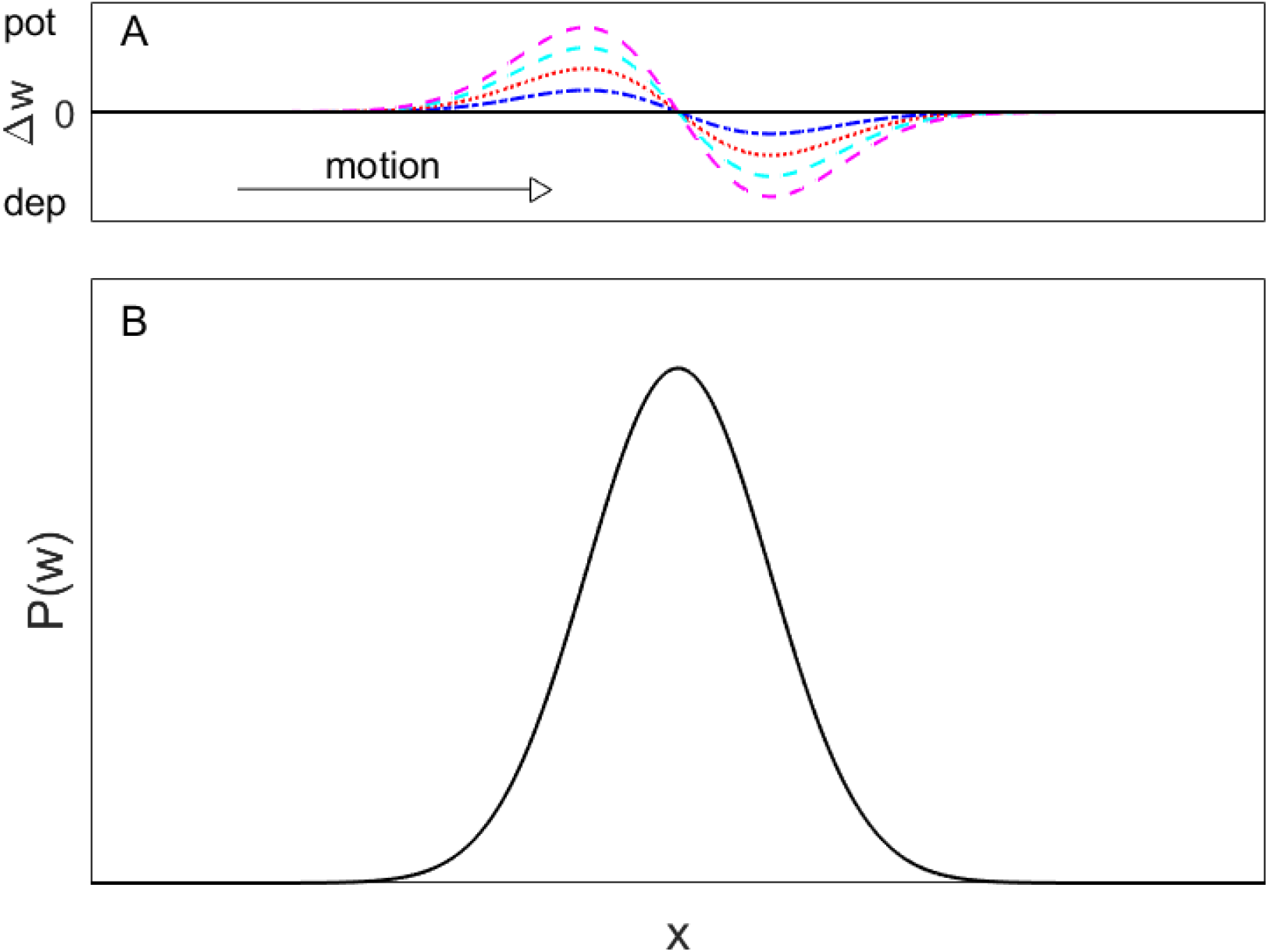
Relative change in weights generated by STDP for motion of the point-stimulus to the right. A) Each plotted line represents the change in weight, Δw, generated by a stimulus with velocity, v = 0.05 (blue dash-dot), 0.1 (red dotted), 0.15 (cyan dashed), 0.2 (magenta solid) cycles/sec, plotted relative to the x-coordinate of the input (pre-synaptic) neuron. The Δw = 0 line is plotted as a black solid line, with potentiation (pot) indicated for Δw > 0 and depression (dep) indicated for Δw < 0. B) The initial Gaussian probability distribution of weights for one post-synaptic neuron in the upper layer neuron receiving pre-synaptic inputs from the lower layer (as shown in Fig.2).

For more realistic two-dimensional motion, as occurs with neural activation of the retina, the above analysis can be extended in a straightforward fashion to two dimensions. Note that this expression does not depend upon the time delay *t*_delay_, which is to be expected since the delay is the same for all neurons between Layers 1 and 2 and it is only the relative times of the pre-synaptic and post-synaptic events at the synapses that play a role in STDP. While this expression for *Δw_ji_* describes the essence of the mechanism by which the shift in weight distribution is in the direction opposite to the direction of motion of the stimulus, there are a number of approximations that will influence the magnitude of this effect. The above calculation uses the continuum expressions for the weight density, whereas in reality each neuron has discrete number of synaptic inputs, which introduces a discretization error to the above calculation. Also, for larger stimuli it is necessary to use the more complete expression, Eq.(6), as this expression takes account of the variability in spiking associated with stimuli with greater spatial extent. In addition, this expression provides a description of only the initial phase of learning, as there are two features of the neural behaviour that make it more difficult to give an analytical expression for the complete weight evolution. First, the weights have an upper bound (i.e., once a weight reach this upper bound, it cannot be further potentiated), which introduces a non-linearity into the weight evolution. Second, the probability of the Level 2 neuron spiking depends upon the complete distribution of all its input weights, which are themselves constantly changing through STDP. In the above expression only the initial Gaussian distribution is used, whereas when learning progresses the distributions will become increasingly non-Gaussian, as described by the receptive fields observed in the numerical simulations in the Results, Section 3.2.

### 2.4 Neural Simulations

Simulations of a network were carried out using MATLAB with *N*_n_ = 2000 neurons at both the input layer and the first layer. The input layer had *N*_p_ = 32 place-fields, i.e., the width *σ*_p_ of the Gaussian place-fields on the input layer was chosen to be *σ*_p_ = 1/*N*_p_ in order to give place-fields that were both localized and had a reasonable amount of overlap. The weight distribution width, *σ*_W_, was also chosen to be *σ*_W_ = 1/*N*_p_, and the width of the resulting place field *σ*_RF,*j*_ of each neuron in the second layer was measured from the distribution of weights after learning by spatially binning the weight amplitudes and finding the width of the resulting histogram, fit to a normal distribution. The width of the place fields in the second layer are distributed around a value that will shift depending upon the velocity of the stimulus, so these are labelled as *σ*_RF,*v*_, to indicate this velocity-dependence. A time-step of 1ms was used for the simulations, and velocities over the range *v* ∈ [0, 5], where the units of velocity are represented in terms of the inverse time (sec^−1^) taken to traverse the full spatial range *x* ∈ [0, 1].

## 3. Results

The analysis here focuses upon the first two layers of the network, since the structure and function of the network follows the same principles at each successive level of the layered network.

### 3.1. Stationary input stimulus

In order to illustrate the effects of a moving stimulus and to have a baseline for comparison, we consider first the case with stationary inputs, i.e., in which the stimulus velocity is zero. We use a single point stimulus with an amplitude 20 times greater than the base firing rate. In this case, there is no change in stimulus position over time, but rather stationary stimuli are presented for short periods of time at random positions. When the stimulus activates the input, it will generate activity in the units at the first layer, as described in Section 2.2 and illustrated in Figure 2. Because this generates constant input to layer two, and a balanced STDP window is used, convolution by the STDP function would not be expected to systematically change synaptic weights. Consequently, the network maintains a stable position code in each layer of the network, namely a localized (Gaussian-like) place field at each layer that arises through the variance of the STDP learning of the weights (Kempter et al., 1999).

The organisation of the receptive fields of the neural populations in the second layer therefore simply reflects the input in the first layer, which has a spatial spread *σ*_p_, and the activity transmitted through the weights, which has a spatial spread of *σ*_W_. Figure 4 shows the results of a simulation for this case, in which the neural population in the second layer generates a place field representation of the input, as expected. The weights are initialized with a small value and then evolve under STDP, as described in the Methods. In this way, the width of the place field distribution at any layer depends upon the width of the place field at the preceding layer and the spatial spread of the synaptic connections that connect the two layers.

**Figure 4:**
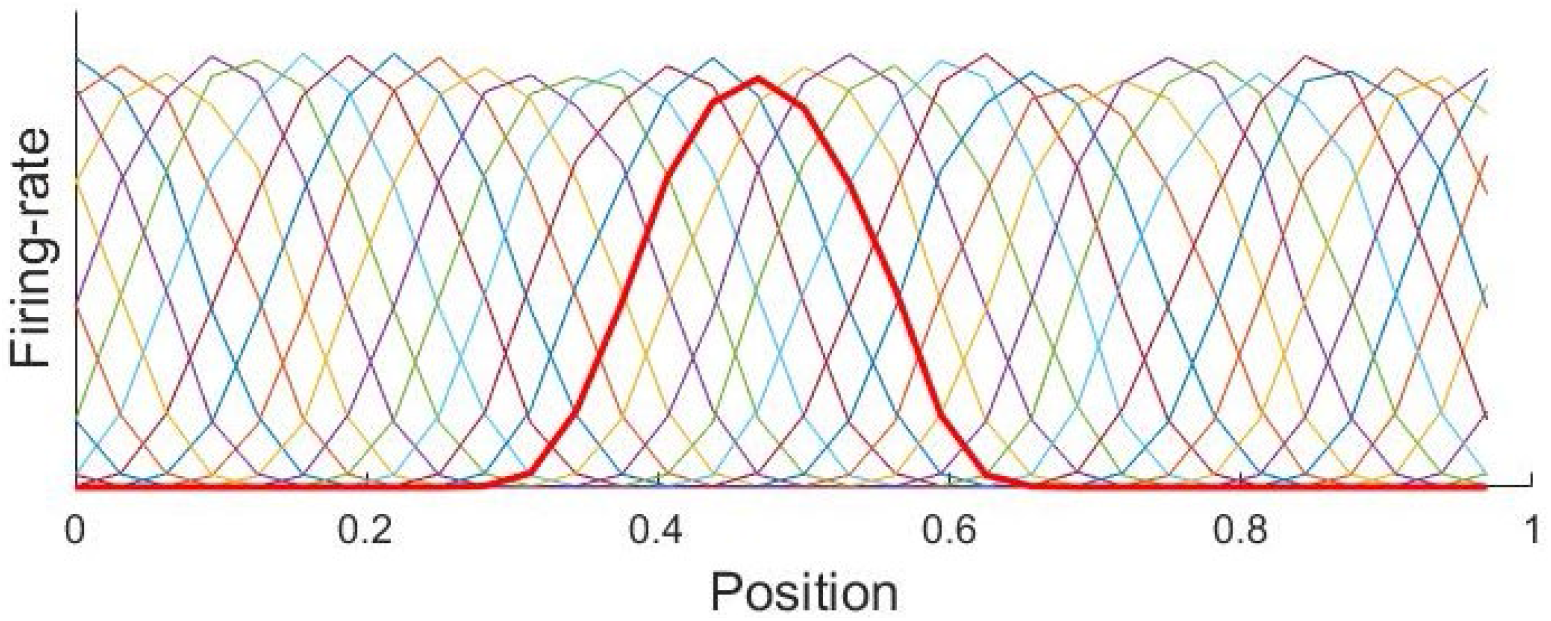
Place code in the second layer generated by learning: Each colour represents the neural activity of a population of neurons, illustrated here for a simulation with N_p_ = 32 position populations in the second layer. The place distribution for the population centred at the position x = 0.5 is shown in bold.

### 3.2. Moving input stimulus

We now consider the case where the same point-stimulus is moving. The velocity is chosen to have a discrete representation (i.e., a discrete number of velocities are chosen for simplicity, rather than a continuous representation), the place input distribution is as described above by a Gaussian distribution of width *σ*_p_, as outlined above, and a simulation time step Δ*t*=1ms is used. A stimulus moving at a velocity *v* has a place representation that changes over time so that at a time Δ*t* later it has shifted a distance Δ*x* = *v* Δ*t*. A moving object will sequentially activate successive populations of Level 1 neurons, which in turn project to Level 2. Importantly, a neuron in level 2 receiving input from level 1 neurons driven by this moving object will tend to fire more as the stimulus moves towards the centre of its place field. Due to STDP, inputs that arrive at the Level 2 neuron relatively early (before its peak firing rate) will be potentiated, whereas inputs that arrive relatively late are likely to be depressed. Consequently, the synapses connecting a given Level 2 neuron to Level 1 neurons centred on the direction from which the stimulus is arriving will tend to be potentiated by STDP. Conversely, the synapses on the other side (i.e., where the stimulus departs from the place field), will tend to be depressed, since the inputs on average arrive after the peak in output spiking activity.

We therefore hypothesise that for neural populations tuned to visual motion, the pattern of arrival of synaptic inputs, together with STDP, will tend to potentiate the synapses in the incoming direction of the stimulus and depress synapses in the departing direction of the stimulus. This would then lead to an overall shift of the place field in the direction towards the incoming stimulus. Moreover, due to the limited temporal window of STDP, the shift in the place fields of the level 2 neurons would be expected to be larger for larger velocities.

To investigate this hypothesis, we simulated the activity of the neural network when it was presented with simulated objects moving at a range of velocities, and investigated the evolution of the receptive fields of Level 2 populations over time. We investigated neural populations tuned to 26 velocities, from 0 to 5 cycles per second in steps of 0.2 cycles per second (since we used periodic boundaries, one cycle is equivalent to traversing the full range of positions once). Because of the symmetry in our neural model, we only considered rightwards velocities, but the network behaves equivalently for leftward velocities. Each simulation ran for 5 simulated seconds (5,000 timesteps of Δ*t* =1ms). The simulated object at a single location provided input to the Level 1 neurons according to their respective place fields.

To evaluate whether receptive fields indeed shifted as a result of learning, we calculated the mean receptive field of all Level 2 neurons at each timestep by aligning the 32 Level 2 place fields and averaging their receptive fields at that timestep. This yielded a mean receptive field as a function of simulation time for each velocity. Figure 5 shows the evolution of receptive field position over time for 6 evenly-spaced velocities (0-5 cycles/s).

**Figure 5:**
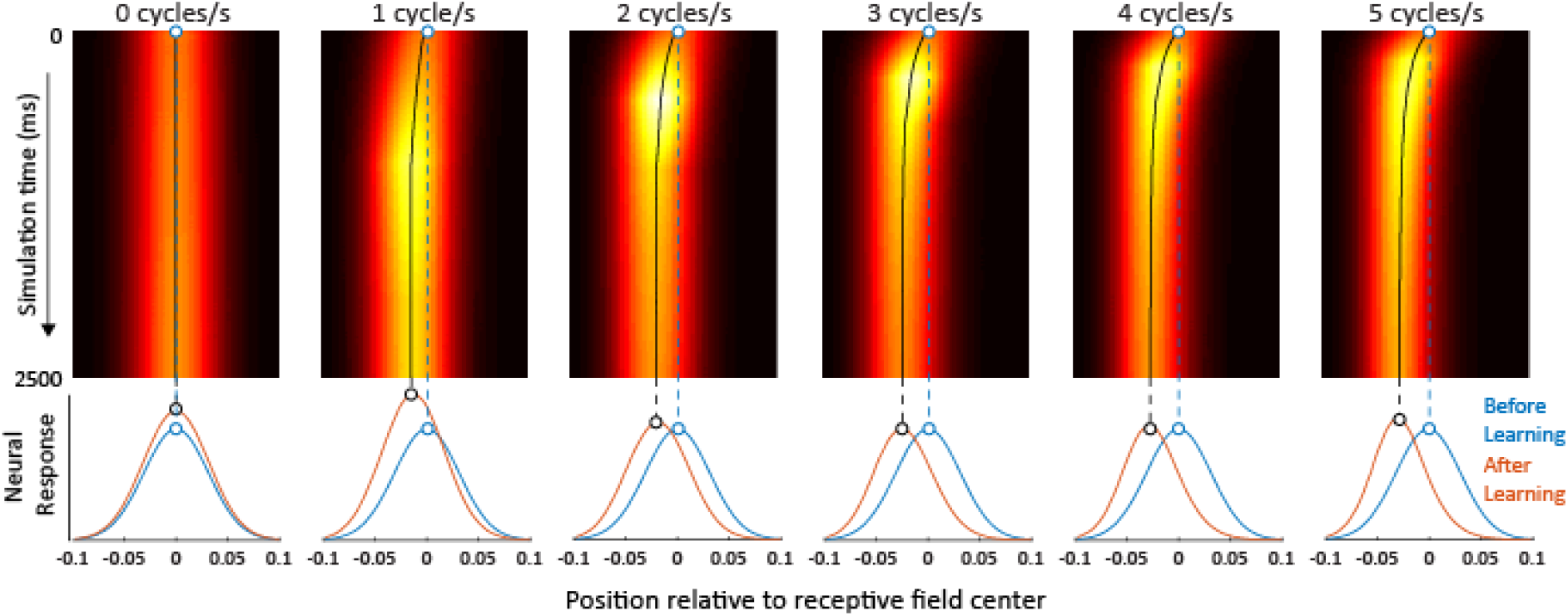
Simulated receptive field position of level 2 neural populations as a function of learning, for 6 different rightward velocities. Top row: In each panel, the colours in each row of pixels show the horizontal receptive field position at a single time-step, with hot colours indicating stronger connections to incoming signals from level 1 populations centred on that position. The center of the receptive field at each timepoint is marked by a black line. For clarity, only the first 2500 ms of the simulation are shown. Bottom row: mean receptive fields before (blue) and after learning (red). Altogether, these plots reveal that when velocity-tuned neural populations are presented with their preferred stimulus (here rightward), STDP causes their receptive field to shift over time in the direction opposite to their preferred velocity (i.e., here leftward).

### 3.3. Velocity-dependence

To be able to directly compare how the evolution of level 2 receptive fields depended on velocity, for each velocity we fitted Gaussians to the average receptive field at each time-step (e.g., each row in each panel in Figure 6). We then repeated the entire simulation 15 times to reduce the impact of stochastic noise. Subsequently, we averaged the horizontal centre of the best-fit Gaussians across all 15 iterations. Finally, we plotted this receptive field centre as a function of time, separately for each velocity. The result is illustrated in Figure 6.

**Figure 6:**
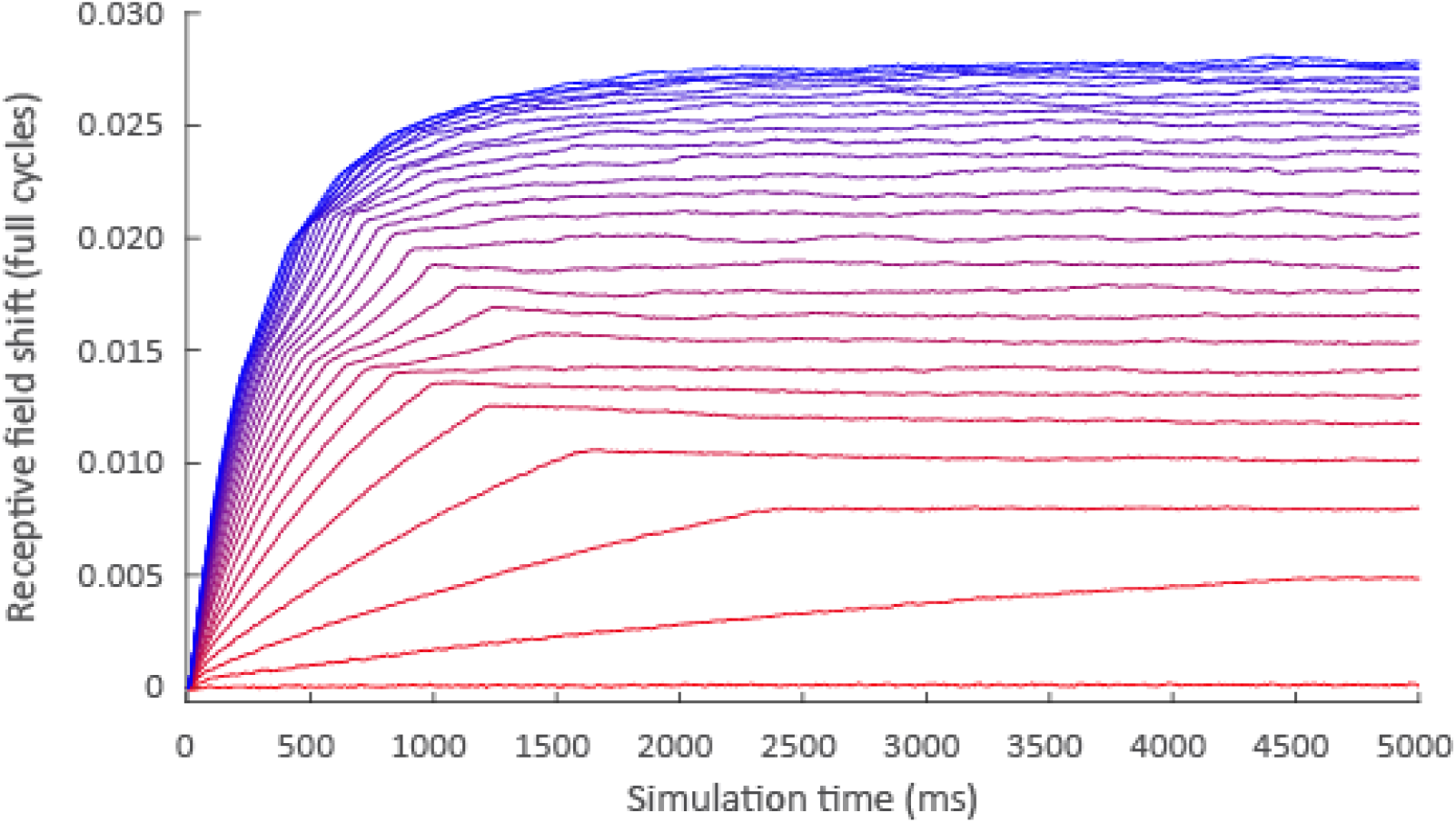
The evolution of receptive field centre over time, as a function of velocity. Colours represent populations tuned to 26 equally-spaced velocities, ranging from zero (red) to 5 cycles/s (blue). For all non-zero velocities, receptive fields shifted against the direction of motion, with a clear monotonic relationship between velocity and the asymptotic magnitude of the shift.

We observed two key features. Firstly, the initial rate at which the receptive field centres of the different velocity-tuned populations shifted increased with velocity. At zero velocity the receptive field centre stayed in the same position, and as velocity increased, the initial rate of change grew until it reached an asymptote. This is perhaps unsurprising, because at lower velocities the simulated object needs more time to traverse the receptive fields of a given number of neurons. As a result, the object drives fewer individual neurons, and in turn provides fewer opportunities for the network to learn.

Furthermore, the neural populations tuned to different velocities differed not only in their initial rate of change, but also in the asymptote of that change. In other words, the spatial shift in receptive field position at which subsequent time-steps produced no further net change in position increased with increasing velocity. This observation is significant because the asymptote represents the position of the receptive field after learning has effectively completed, and therefore reflects the stable situation in visual systems that have had even a short history of exposure to moving stimuli. The rate at which the asymptote is approached depends upon the STDP learning rate. the Because these velocity-tuned populations are subpopulations of an overall population coding for position (as illustrated schematically in Figure 2), the overall population effectively represents a moving object ahead of where a physically-aligned static stimulus is represented. As a consequence, we might expect the asymptotic receptive field shift to be similarly reflected in conscious perception as the instantaneous perceived position of a moving object.

### 3.4 Behavioural predictions

Our model reveals how STDP-induced shifts in receptive field position depend on velocity. In the final section of this paper, we evaluate the degree to which these predictions match observed dependencies on velocities in the localisation of moving objects by healthy human observers.

A much-studied behavioural paradigm used to probe the instantaneous perceived position of a moving object is the flash-lag paradigm, in which a flashed object is briefly presented alongside a moving object (Nijhawan, 1994). Observers are then required to report where they perceived the moving object to be at the moment the flashed object was presented. Strikingly, in this paradigm observers consistently localise the moving object ahead of the physically aligned flashed object, a phenomenon known as the Flash-Lag Effect (FLE) (Nijhawan, 1994). Although the mechanisms underlying this effect have been hotly debated over the last 25 years, convergent evidence supports Nijhawan’s original proposal that it reflects some kind of neural motion extrapolation process (Hogendoorn, 2020; Nijhawan, 1994). What is particularly relevant to the present context is that the effect has been observed to scale with velocity (Wojtach et al., 2008): when an object moves faster, its perceived position at any given instant lies further along the object’s trajectory.

In our model, the perceived position of a moving object corresponds to the level 2 neural population activated by that object. As outlined in the previous section, this is determined by the asymptotic receptive field position after learning. As a measure of asymptotic receptive field shift after learning, we averaged the receptive field shift in the final 100 ms of our simulation (e.g., the 100 rightmost datapoints for each curve in Figure 6), averaged across the 15 iterations of the simulation. We subsequently fitted a logarithmic function to the data, as has previously been done for behavioural estimates of perceived position shifts using the FLE (Wojtach et al., 2008). This function explained a total of 96.8% of the variance, showing that the dependence of final receptive field position on velocity was very well described by a logarithmic relationship (Figure 7).

**Figure 7:**
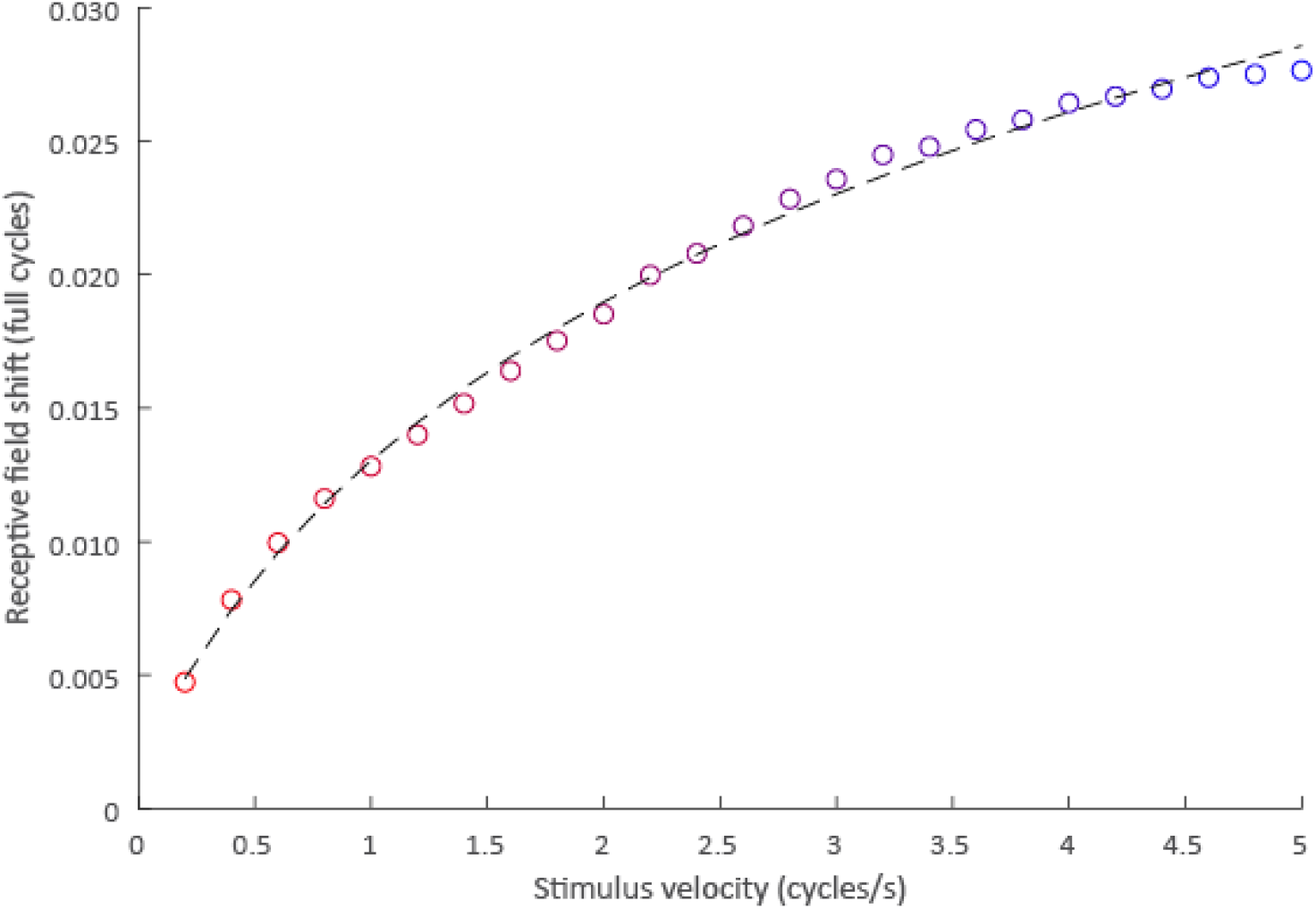
Velocity-dependence of asymptotic receptive field shifts. Marker colours correspond to velocities in Figure 6, and the dashed line represents a logarithmic fit to the data, explaining 96.8% of variance.

In order to compare the perceptual shifts predicted by our model to those measured in behavioural experiments with human observers, we compared the velocity dependence of receptive field shifts in our model to the velocity dependence of the FLE, as previously measured for the full range of detectable velocities by (Wojtach et al., 2008). We noted that the magnitudes of both RF-shifts in our model and perceptual shifts in the FLE were very well-described by a logarithmic dependence on stimulus velocity (Figure 7, Figure 8A). We then directly compared RF-shifts in our model to perceptual shifts in the FLE by treating the maximum velocities tested in each paradigm to be equal. For the behavioural paradigm, this was 50 deg/s, the highest velocity at which an FLE could be measured (Wojtach et al., 2008). For our model, this was 5 cycles/second, at which point the period of the motion (200 ms) reached the approximate width of the STDP window. The correlation between RF-shifts in our model and perceived position shifts in the FLE was near perfect (r > 0.99). Note that this pattern of results arose spontaneously as a result of STPD, without requiring any tuning of the model. This shows that the velocity-dependence of STDP-induced receptive-field shifts in our model very closely matches the velocity-dependence of perceptual mislocalisation for moving objects as measured using the FLE.

**Figure 8:**
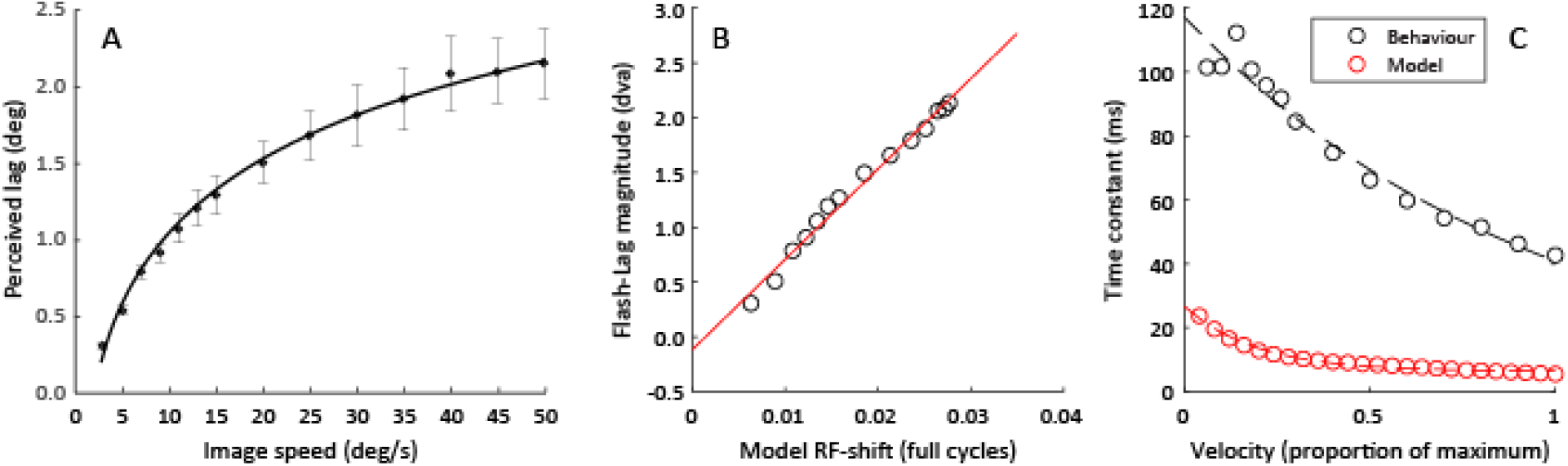
A) Behaviourally measured perceptual shift magnitude as a function of velocity using the flash-lag effect (reproduced from (Wojtach et al., 2008). The solid line indicates a logarithmic fit to their data. B) Scatterplot showing the magnitude of receptive field shifts predicted by our model and the corresponding perceptual shifts reported by (Wojtach et al., 2008) using the FLE. The correspondence is near-perfect, with a Pearson correlation r > 0.99. C) Absolute magnitudes of RF-shifts produced by our model and position shifts observed in the FLE, expressed as a time-constant (the equivalent duration necessary to traverse the shift distance at each velocity). Dashed lines indicate exponential fits.

Finally, we compared the absolute magnitude of the shifts in receptive field position produced by our model to the absolute magnitude of perceptual shifts observed in the FLE. To do so, we expressed the magnitude of the shift at each velocity as a time constant, by dividing shift magnitude by velocity. This is equivalent to the time necessary for an object moving at that velocity to be displaced a distance equivalent to the receptive field shift (Figure 8C). We observed that for both the FLE and our model, this time constant tended to decrease exponentially with increasing velocity (exponential fits explained 96.3% and 98.5% of variance in FLE and model time constants, respectively). Across the entire range of velocities tested, the time constant produced by our model was roughly 12-20% of the time constant for the behaviourally-measured FLE, as might be expected given that our model reflected receptive field shifts in just a single layer of synaptic connections.

### 3.5 Parameter dependence

Parameters in our model were chosen to be biologically plausible. For some parameters, choosing different values would be expected to have predictable effects. For example, varying the STDP learning rate *ρ*, Eq.(2), would be expected to cause the model to converge to its asymptotic state either more rapidly or more slowly. However, we would not expect it to change the asymptotic state itself – merely the simulation time necessary to reach that state. Indeed, we deliberately chose a relatively high learning rate in order to keep the computation tractable; we would not expect a biological system to reach its asymptotic state within just 5 seconds of exposure.

For other parameters, it is less obvious how choosing different values would affect the pattern of results. In particular, we chose a value of 32 for Np, the number of place fields in each layer. This parameter corresponds loosely to the size of receptive fields at each layer, and might be expected to vary for neurons in different areas in retinotopic visual cortex. For example, place field width inevitably varies as a function of eccentricity, with foveal retinotopic areas showing smaller receptive fields than peripheral retinotopic areas. To investigate the effect of manipulating this parameter, we ran additional simulations with higher (64) and lower (16) values of Np. Although we observed small differences in the absolute magnitude of predicted receptive field shifts, the overall pattern of results was very similar (Figure 9). In particular, the pattern of velocity dependence for both absolute receptive field shifts and the equivalent time constants was highly similar, giving confidence that our results are not restricted to a small region of parameter space.

**Figure 9:**
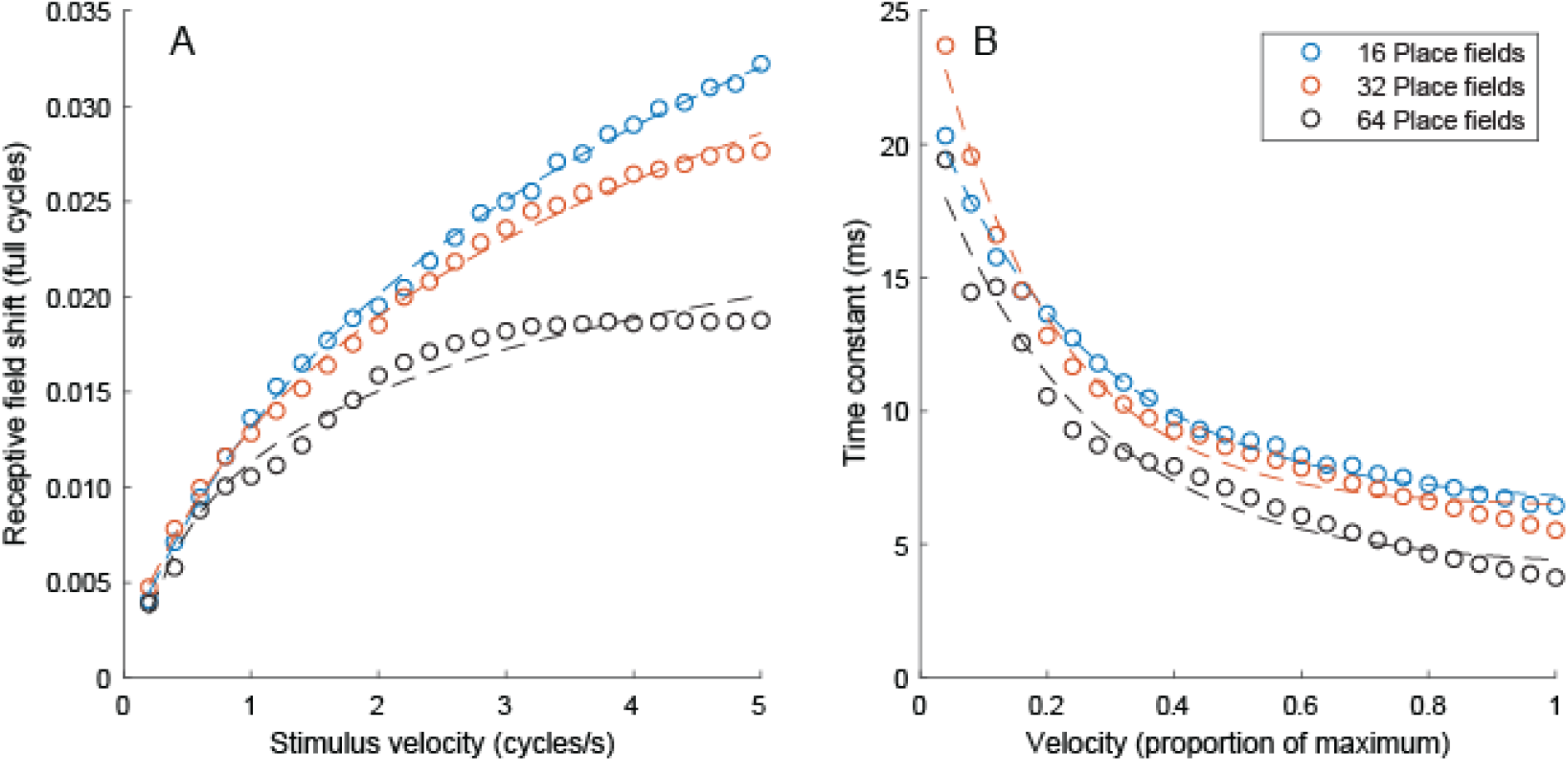
Comparison of model prediction for different numbers of place fields. A) Receptive field shift as a function of velocity for models with 16, 32, or 64 place fields. The magnitude of the receptive field shift increases slightly with increasing place field width (corresponding to a lower number of place fields), but the overall pattern of velocity-dependence is comparable for all three simulations. Dashed lines show logarithmic fits as in the main analysis. B) Comparison of time constants for models with 16, 32, or 64 place fields. Time constants were remarkably similar for models with different numbers of place fields, showing comparable exponential dependence on velocity. Dashed lines show exponential fits.

## 4. Discussion

We investigated a computational problem faced by the brain in processing the visual position of moving objects: the fact that neural transmission takes time, and that the brain therefore only has access to outdated visual information. Motion extrapolation is one way the brain might compensate for these delays: by extrapolating the position of moving stimuli along their trajectory, their perceived position would be closer to the their physical position in the world (Hogendoorn, 2020; Nijhawan, 1994, 2008; Hubbard, 2005). We simulated a possible neural mechanism (spike-timing dependent plasticity - STDP) by which a layered neural network might implement such an extrapolation mechanism. We show that a two-layer hierarchical network comprised of velocity-tuned neural population is not only able to implement motion extrapolation, but actually learns to do so spontaneously without supervision, due only to the temporal contingencies of its connectivity. We go on to show that the velocity-dependence of the resulting receptive field shifts predicts previously reported, behaviourally measured effects on the perceived position of moving objects.

The magnitude of the receptive field shifts we observe for each velocity in our simulations corresponds roughly to the equivalent displacement resulting from 10-20 ms of motion at that velocity. Although this is 5-8 times smaller than the FLE, it is important to note that our model analysed here includes just two layers, and only one stage at which learning takes place. If we were to extend our model to include additional layers, each with comparable properties, then each output layer would constitute the input layer for the synapses at the next stage. As a result, we would expect the same learning process, and therefore the same receptive field shift, to take place at each stage. In this way, receptive field shifts would add up as information ascends the hierarchy. Although it is unknown which cortical areas ultimately determine where we consciously perceive a moving object, it seems highly likely that information from the retina will cross at least a handful of synapses before it is accessed for conscious awareness. The magnitude of the receptive field shifts predicted by our model are therefore of roughly the same order of magnitude as, and comparable to, those we would expect based on the magnitude of the perceptual effect.

It is interesting to note that the flash-lag effect is just one of several related motion-position illusions in which the position of a moving object is biased by motion (e.g., Eagleman & Sejnowski, 2007). In the Fröhlich effect (Kirschfeld & Kammer, 1999) for example, a moving object suddenly appears, and the perceived initial position of the object is mislocalised in the direction of motion. The pattern of receptive fields shifts that we observe in our model can also qualitatively explain the Fröhlich effect. Subtle differences with the FLE paradigm (such as the likely transient neuronal onset response to the initial appearance of the moving object in the Fröhlich effect) are not currently captured by our model, but a direct comparison of predictions for these (and other) illusions could be informative to further develop the model.

The magnitude of receptive field shifts predicted by our model is also consistent with previous neurophysiological recordings as well as human neuroimaging. Jancke and colleagues (Jancke et al., 2004) recorded neurons in cat primary visual cortex, and compared the latencies of responses to flashes with the latencies of responses to smoothly moving objects. They observed that peak neural responses to smoothly moving objects were approximately 16 ms further along the motion trajectory than peak responses to static flashed objects. Almost identical results were found by Subramaniyan and colleagues (Subramaniyan et al., 2018) who recorded neurons in primary visual cortex of awake macaques, and observed a latency advantage for moving stimuli compared to flashed stimuli of between 10-20 ms depending on stimulus velocity. These results from invasive recordings in cats and macaques are therefore in quantitative agreement with the predictions of our model. In humans, we recently used an EEG decoding paradigm to investigate the latency of neural responses to predictably and unpredictably moving objects (Hogendoorn & Burkitt, 2018). Using an apparent motion paradigm, we showed that when objects move along predictable trajectories, their position is represented with a lower latency than when they move along unpredictable trajectories. Like the neurophysiology studies, we observed a latency of 16 ms for the predictably moving object. Our present modelling result is therefore consistent not only with behavioural measurements of motion perception, but also with neural recordings in both humans and animals.

The underlying mechanism is the same property of STDP that causes neurons to tune to the earliest spikes (Song, Miller & Abbott, 2000; Guyonneau, Van Rullen & Thorpe, 2005), which has diverse manifestations in the brain’s neural circuits, including in the context of phase precession in the hippocampus (Mehta, Quirk & Wilson, 2000) and of localizing a repeating spatio-temporal spike pattern embedded in a noisy spike train (Masquelier, Guyonneau & Thorpe, 2008). Essentially STDP leads to potentiation of the weights associated with the leading edge of the receptive field as the receptive field is entered, and depression of weights on the tailing edge as the motion moves out of the neuron’s receptive field as illustrated in Figure 3. Our study extends the understanding of this mechanism by examining the velocity dependence of this effect. It is important to note that in our model, the extrapolation mechanism emerged spontaneously and without supervision, simply as a result of STDP. By extension, extrapolation would similarly be expected to develop spontaneously in *any* hierarchical network of velocity-selective populations when it is exposed to visual motion. Furthermore, it would be expected to arise between every layer in such a network. This structure of extrapolation mechanisms at multiple levels of the visual hierarchy is consistent with previous empirical findings showing extrapolation at both monocular and binocular stages of processing (van Heusden et al., 2019). It is also consistent with the Real Time Temporal Alignment hypothesis that we recently proposed (Hogendoorn and Burkitt, 2019) as a theoretical extension of classical predictive coding (Rao & Ballard, 1999), although this hypothesis also posited feedback projections that are not included in our current model. Indeed, the network architecture considered here is entirely feed-forward, which represents a good model for the initial wave of neural activity travelling through the visual pathway in response to a stimulus, but it neglects the feedback activity from higher visual centres. This descending feedback activity, which occurs and can persist over a longer timeframe than the initial wave of neural activity, may play an important role in the understanding of temporal processing in the brain on these longer timescales. These questions, which lie outside the scope of the present study, form the focus of ongoing research.

The mechanism here differs from that proposed by Lim & Choe (2006), who show that STDP, together with facilitating synapses, can provide a neural basis for understanding the orientation flash-lag effect, a visual illusion involving the perceived misalignment of a rotating bar, which is located between two aligned flanking bars that briefly flash when the rotating bar is aligned. Importantly, their model architecture is very different, involving a bilaterally ring-connected network of orientation-tuned neurons, in which the lateral connections between these neurons are trained using a combination of STDP and activity-dependent facilitation, rather than the feedforward connections examined in our study. They also did not examine the velocity dependence of the effect.

The network and learning parameter values chosen for this proof-of-concept study represent values consistent with cortical neural processing, but without incorporating many of the details of vision processing in the human visual pathway. The description of the neural activity in terms of a Poisson process is a widely used approximation for the time distribution of action potentials. Although it neglects all spike after-effects, such as refractoriness, it nevertheless provides a good description for the situation examined here in which the visual stimulus moves with a constant velocity and is modelled as having a spatial intensity distribution that is Gaussian, without any edges or other spatial discontinuities (Aviel & Gerstner, 2006). The STDP time constants of potentiation and depression, *τ*_p_ and *τ*_d_, are chosen to both have a value of 20ms, which is in the range of that observed in neurons in the visual cortex (Froemke & Dan, 2002).

In the primate visual system, the processing of motion begins in the retina, where it has long been known that there are direction-selective retinal ganglion cells (Barlow & Hill, 1963; Barlow et al., 1964). These neurons are maximally activated by motion in their preferred direction and strongly suppressed by *motion in the opposite direction.* Within the retina there are a number of mechanisms involving multilayered retinal circuits that provide reliable motion detection (Frechette et al., 2005; Manookin et al., 2018), consistent with the broad experimental evidence for velocity tuning neurons in the retina of mammals and vertebrates (Olveczky et al., 2007; Vaney et al., 2012; Ravello et al., 2019; for a review see Wei, 2018). This motion selective information is transmitted via the LGN to the primary visual cortex, V1, where direction selective neurons are concentrated in layer 4B of V1 and project from there to higher motion processing areas of the visual hierarchy pathway, particularly area MT (Maunsell & van Essen, 1983).

In the analysis presented here we have made the simplifying assumption that the same principles apply at each successive level of a layered network. However, in the visual system the receptive fields of neurons at successive levels become progressively larger as information moves up the visual hierarchy. For example, a motion selective neuron in area MT which has a receptive field of 10^0^ diameter receives its input from neurons in V1 that have receptive fields of 1^0^ diameter (Andersen, 1997). The restricted receptive field of neurons in the lower stages of this vision processing hierarchy can result in ambiguous motion signals as a result of the aperture effect. Consequently, at each successive stage of the visual hierarchy the motion information is not only transmitted, but it also can be refined: the information at earlier stages is integrated so that the motion of larger objects can be more accurately determined and objects moving at different speeds are disambiguated – for a review see Bradley & Goyal (2008). The larger receptive field sizes in the higher stages of the hierarchy correspond to broader place field representations, which it would be straightforward to accommodate in a multi-layer extension of the processing framework presented here. It may also be possible that the dorsal and ventral pathways of the visual system (i.e., the where and the what pathways) have very different encoding of velocity. While the dorsal pathway relies upon an accurate representation of position and the visual motion extrapolation analysed here, it is possible that in the ventral pathway the velocity coding is so broadly tuned that it is effectively absent.

In the analysis presented here we have for convenience used a discrete coding of the velocity, rather than allowing it to take a continuum of values from zero up to some maximal value. Consequently, an object with changing velocity will, in this simplified model, make discrete jumps between velocity sub-populations. It is, however, also possible to formulate the velocity using such a continuous representation, for example as a set of overlapping Gaussian distributed velocity fields similar to the (spatial) place fields. We anticipate that this would give a smoother, possibly more biologically plausible, transition between velocity sub-populations, but that it would not change the essential results of this study in any significant way.

In sum, we have implemented spike-time dependent plasticity in a layered network of velocity-selective neurons, and shown that this results in a pattern of receptive-field shifts that causes the network to effectively extrapolate the position of a moving object along its trajectory. The magnitude of this shift is in quantitative agreement with previous findings from both animal neurophysiology and human neuroimaging experiments, and also qualitatively predicts the perceptual mislocalisation of moving objects in the well-known FLE. Most strikingly, we show that it emerges spontaneously and without supervision, suggesting that extrapolation mechanisms are likely to arise in many locations and at many levels in the visual system.

Finally, the model we present here includes only feed-forward connections, and a natural extension to the model would be to include lateral and/or feedback connection. Previous modelling work, most notably by Jancke and Erlhagen (Jancke & Erlhagen, 2010), has proposed an instrumental role for lateral connections in generating the perceptual mislocalisation that characterises the FLE. It would be interesting to investigate in more detail what the emergent characteristics would be of a network implementing both STDP and lateral connectivity, and whether that would explain any other perceptual phenomenology. In a similar vein, it would be interesting to implement feedback connections in the model we present here, as we proposed in our previous Real-Time Temporal Alignment hypothesis (Hogendoorn and Burkitt, 2019). An exciting possibility is that these feedback connections might function to calibrate receptive field shifts to the relative transmission delays between layers in the hierarchy, allowing extrapolation mechanisms to accurately compensate for processing delays. Further research will be necessary to evaluate these possible extensions to the current model.

## Acknowledgments

HH acknowledges support from the Australian Research Council’s Discovery Projects funding scheme project DP180102268. ANB acknowledges funding from the Australian Government, via grant AUSMURIB000001 associated with ONR MURI grant N00014-19-1-2571. We thank Hamish Meffin and Stefan Bode for helpful comments on the manuscript.

